# Qualitative Optimization of Oncolytic Virotherapy and Immune Therapy Combination Treatments

**DOI:** 10.1101/2025.09.15.676375

**Authors:** Negar Mohammadnejad, Thomas Hillen

## Abstract

Oncolytic viruses (OVs) are designed to selectively target and destroy cancer cells while sparing normal, healthy tissue. Several viruses for oncolytic virotherapy are currently developed. In this paper, we will use mathematical modeling to consider key strategies that can improve the efficacy of oncolytic virotherapy. These include the integration of immunotherapy approaches with virotherapy to amplify anti-tumor immune responses, as well as optimizing the timing, dosage, and sequencing of viral administrations. Specifically, we consider strategies that increase the burst size of the virus, immunostimulation and immunosuppression, we optimize for different weekly virus injection schedules, and we consider the combination of OV therapy with CAR-T cell therapy. A limiting factor is the availability of data. We parametrize the model from several different data sets. These, however, relate to different cancers and different experimental set up. Hence our model cannot be considered to be validated. Consequently, our results are qualitative. Our results highlight the critical importance of timing for virotherapy’s efficacy and overall success. They outline strong evidence for promising treatment scenarios that need to be further tested experimentally in the future.

## 1 Introduction

The mechanism of oncolytic viruses (OVs) is similar to that of conventional viruses; they enter the host cells and exploit the cell’s genetic machinery to replicate and spread [14, 28, 32]. What distinguishes OVs is their capacity to selectively target and destroy cancer cells while sparing normal, healthy tissue. The immune system was once considered a limiting factor, with the belief that its response to the virus would inhibit replication, restrict viral spread, and reduce tumor cell killing. However, this view has largely shifted, as it is now recognized that the immune system plays a pivotal role in OV–cancer interactions [17, 33]. Viral infection of the tumor enhances its visibility to the immune system, promoting recognition and attack [17, 25, 30]. Moreover, virus-induced lysis of tumor cells leads to the release of tumor antigens into a microenvironment modified by infection, thus reversing tumor-induced immune suppression. The first genetically engineered OV, a herpes simplex virus 1 (HSV-1) mutant with deficient thymidine kinase, was developed in 1991 and used to treat malignant glioma in nude mice [31]. These viruses preferentially target and eliminate cancer cells, often leveraging dendritic cells to mediate their effects [25].

Baadbdullah et al. [4] extensively studied a mathematical model for the temporal and spatial interactions of the virus and the cancer cells without considering the effect of the immune system. In the absence of OVs, tumors are known to create an immunosuppressive environment, effectively silencing the body’s immune response against cancer cells. However, once oncolytic viruses are introduced, they induce pro-inflammatory signalling within the tumor microenvironment. This shift not only weakens the tumor’s defenses but also enhances its susceptibility to immunotherapies. OVT awakens the immune system, enabling a robust response against the tumor. Therefore, incorporating the immune system into the model is essential. It is important to note that the oncolytic virus may be cleared from the body before treatment can fully eliminate the tumor. Despite the fact that OVs were translated into clinical trials, with T-VECs receiving FDA approval for melanoma, there are still challenges, such as limited viral infection and immune clearance, that need to be overcome [30].

Based on the model of Al-Tuwairqi et al. [2], we incorporate the population of immune cells into a model and build a base model which investigates the fundamental interactions of immune response with cancer and with the oncolytic virus. In [2], the authors conducted a qualitative analysis of the model and its steady states. They examined the existence and stability of equilibrium points, and through numerical analysis they found oscillatory behaviour. Moreover, they demonstrate the impact of various parameters on treatment outcomes. We continue this research by including specific OV treatment strategies and combination treatments. Specifically, we analyze the outcome of an increased effective viral production rate, the effect of immuno-suppression and immuno-stimulation, we optimize the scheduling of OV injections, and we combine OV therapy with CAR-T cell therapy.

We find an important aspect of combination therapies, which is often forgotten in theoretical or experimental studies, is the timing of the treatments. Using our oncolytic virus model, we show that the efficacy of treatment can be significantly boosted compared to the traditional dosing, when a massively different timing is applied. In our case, an optimal schedule gives an initial dose on day one, and additional, larger doses in the following weeks. We obtained that the outcome of the treatment is significantly more successful when the virotherapy is re-administered three or four weeks after treatment began. The sensitive dependence of the treatment outcome on the scheduling is surprising, and offers a simple way to improve these treatments.

After exploring various treatment variations, we extended our analysis of the OVT base models by incorporating a more specific representation of immune responses and analyzing the same treatments within this enhanced framework. We find essentially the same results as the more complex model. Also in this case, a careful treatment scheduling of viral injections makes a huge difference in the model outcome.

It needs to be pointed out that our work has not be validated against data and results that we have obtained in this work should not be considered as treatment recommendations. Our results are quantitative. They outline strong evidence for promising treatment scenarios that need to be further tested experimentally in the future.

### 1.1 Previous Modeling of OV Therapy

There is a considerable history of mathematical modeling of oncolytic virotherapy. The works by Baabdulla et al. [4] and Al-Tuwairqi et al. [2], mentioned earlier, build on this foundation. One of the earliest models was introduced in 2001 by Wodarz [45]. His proposed model explored how virotherapy could be optimized to achieve maximal therapeutic effects. Wodarz examined three main mechanisms of action: direct viral killing of tumor cells, immune responses against the virus that indirectly aid therapy, and the induction of tumor-specific immune responses triggered by the infection. The analysis demonstrated that simply increasing the virus’s cytotoxicity is not always beneficial, as overly rapid killing can limit virus spread and reduce treatment efficacy. Instead, an optimal balance must be struck between viral replication and cytotoxicity. Building on this, in 2009 Komarova and Wodarz [10] presented a broader modeling framework that accounted for different modes of virus spread (fast vs. slow) and tumor growth kinetics. Their key insight was that the dynamics of OV therapy can fall into two qualitatively distinct regimes, the fast spread and the slow spread. In the following year, in [27], they continued their study of the OVT dynamics and developed a general system of ODEs that builds on biologically motivated properties of tumor growth and viral spread. They analyzed how varying these assumptions would alter system dynamics and defined conditions under which virus-mediated tumor eradication would be possible. Their work also underscored the importance of model robustness and cautioned that many classical modeling results may be artifacts of arbitrary mathematical choices. Their work on oncolytic virotherapy has served as a foundational framework upon which more complex models has been constructed. In 2011, Tian [42] introduced a model comprising of three variables, the uninfected and infected cancer cells, and the virus and his analysis showed the burst size of the oncolytic virus plays a crucial role in the success of virotherapy. His findings underscored the importance of optimizing the burst size of oncolytic viruses in virotherapy. In the other studies of the OVT, the focus was on the role of the immune system in the cancer-virus interactions. In the work done by Eftimie et al [11], a detailed ODE-based model with two compartments (lymphoid and peripheral) was developed to represent interactions between uninfected/infected tumor cells, memory/effector immune cells, and the two viruses-adenovirus (Ad) vaccine and an oncolytic vesicular stomatitis virus (VSV). This model could accurately reproduces the experimental tumor dynamics and immune responses. Using this mathematical model, they explored the conditions necessary for achieving permanent tumor elimination. They had their focus on two complex dynamic behaviours: multi-stability and multi-instability. Their work also identified that the prolonged oncolytic virus presence is lined to a distinct phenomenon—multi-instability. Additionally, they assessed whether viral persistence alone was sufficient for tumor elimination. The findings revealed that without an active anti-tumor immune response, even a sustained anti-viral effect could not prevent tumor growth. Thus, successful treatment requires the combined action of both immune-mediated tumor killing and oncolytic viral activity. In another study on the immune-virus interactions Storey et al. [41] distinguished between the innate and adaptive (tumour and virus specific) immune response. They pointed out that the innate immune system clears the virus too quickly, reducing its ability to infect tumor cells and stimulate an effective anti-tumor immune response and as a result, OVT alone is often insufficient for tumor clearance. They modeled a combination therapy of OVT and the PD-1/PD-L1 checkpoint inhibitor. This combination therapy prevents T-cell exhaustion, improves immune-mediated tumor killing, and significantly lowers the viral infectivity threshold required for successful treatment. They highlighted the importance of careful scheduling and dosing in combination therapy. Their analysis pointed out that administering a second viral dose too soon after the first can worsen treatment outcome due to immune system redirection toward viral rather than tumor antigens. In 2021, Pooladvand et al. [36] developed a 3D spatio-temporal model to study the dynamics of cancer-virus interactions, specifically focusing on adenovirus therapy within a solid tumor. In their study, by linking PDE modeling with bifurcation analysis, they revealed deep insights into why virotherapy often falls short as a standalone treatment, and underlined the limitations of virotherapy as a monotherapy. They also highlighted the need for combination strategies to achieve complete tumor control.

## 2 The Mathematical Model

Our model for immune response to oncolytic virotherapy is based on a standard oncolytic virus model as analyzed recently in Baabdullah [4]. This model is well known in the literature and has been used in many publications [45, 42, 36, 2]. The basic OV model consists of a system of reaction-diffusion equations on a domain Ω, and is given by

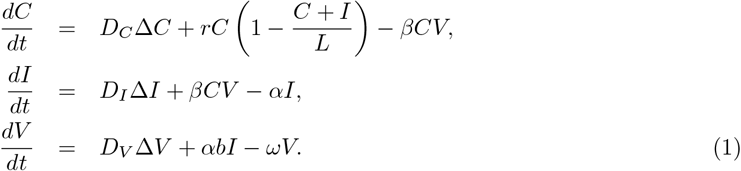

where *C*(*x, t*), *I*(*x, t*), and *V* (*x, t*) represent the populations of uninfected cancer cells, infected cancer cells, and free virus particles, respectively. Their corresponding diffusion coefficients are denoted by *D*_*C*_, *D*_*I*_, and *D*_*V*_ . We have the Laplace operator Δ as the sum of all second order derivatives. The second term in equation (1) describes the logistic growth of cancer cells in the absence of therapy at the rate *r* with carrying capacity *L*. The interactions between tumor cells and virus particles are modeled by the mass action term *βCV* where cancer cells become infected by the virus at a rate *β*. The second term in Equation (1) accounts for the increase in infected cells. The lysis of these infected cells is represented by the third term at the rate *α*. In the last equation, *b* represents the burst size of the virus in infected cells, while *ω* denotes the viral clearance rate.

Model (1) can be considered with standard boundary conditions, for example homogeneous Dirichlet boundary conditions in case of a diffusive cancer, or homogeneous Neumann boundary conditions in case of an encapsulated cancer.

This model (with homogeneous Neumann boundary conditions) was recently analyzed in [4] for pattern formation. In the absence of spatial terms, the above model (1) shows a Hopf bifurcation for large enough burst size parameter *b*. If taken into the spatial context, we then have local oscillators that are coupled through diffusion, and as shown in [4] very interesting and complex spatio-temporal patterns arise. Here, we extend the above model (1), to include immune response, and we focus on the kinetic terms, i.e. we remove spatial dependence for now.

### 2.1 The Basic OV-immune Model

The simplest way to include an immune response into the above model is to add a population of effector immune cells *Y* (*t*) into the model. This was done by Al-Tuwairqi et al. [2], and we will make two small modifications to Al-Tuwairqi’s model. Our very first (space independent) model becomes

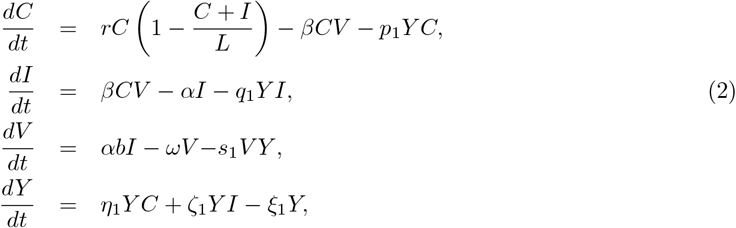

where *Y* (*t*) represent the populations of cytotoxic immune cells. The effect of the immune response on cancer cells, infected cancer cells, and virus is modelled through standard mass action terms with rates *p*_1_, *q*_1_, and *s*_1_, respectively. The last equation of this system describes the stimulation of the immune system by both uninfected and infected cancer cells with rates *η*_1_ and *ζ*_1_, respectively. The immune cells are cleared at rate *ξ*_1_ as denoted in the last term. We will be referring to system (2) as our base model from now on.

In contrast, the model studied by Al-Tuwairqi et al. [2], included a viral loss term of the form *−β*_2_*CV* in the *V* equation. It was argued by many researchers [45, 11, 46], and also in [4] that such a term can be neglected, as a viral infection has a minimal loss effect on the large virus population. Hence, here we choose not to include such a term. Additionally, we choose not to include a term for the natural death of cancer cells, *−dC* in the *C* equation, since on the time scale of OV therapy (few weeks), such a term is not relevant.

In Figures 1a and 1b, we sketch the interactions of the elements of the Baadbullah et al. model (1), and the base model (2) respectively. We consider the system (2) with the initial conditions:

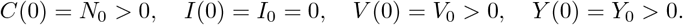

**Figure 1:**
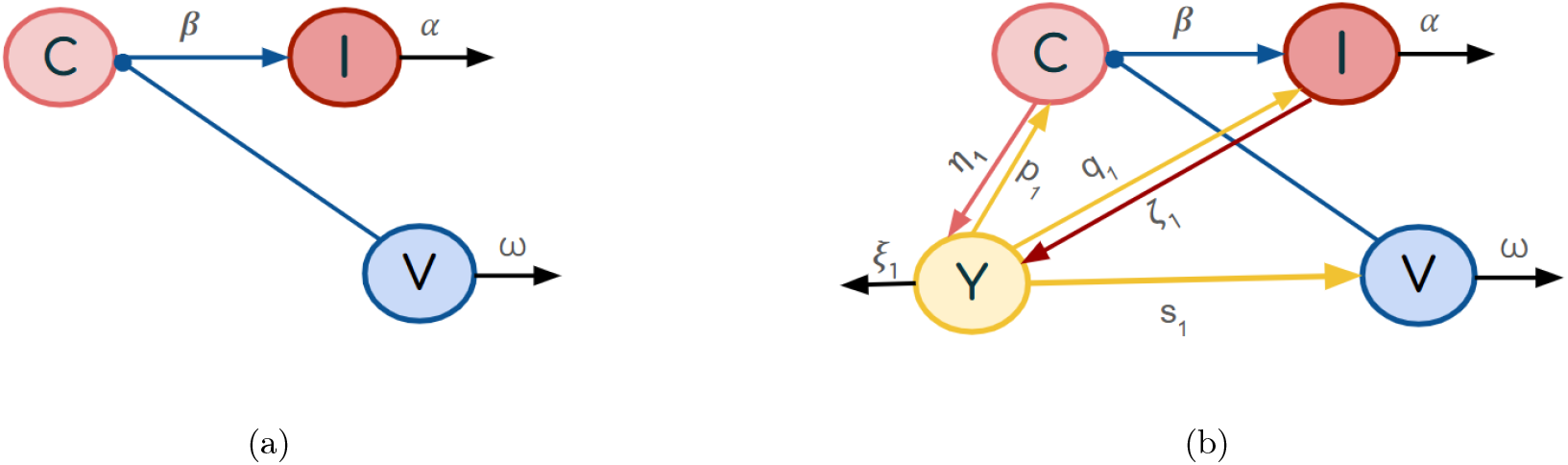
(a) The blue arrows represent the viral infection of cancer cells at rate *β*. All black arrows depict cell death and virus degradation (b) The blue and black arrows represent similar dynamics as in (a). The yellow arrows indicate cell and virus death by the immune system at rates *p*_1_, *q*_1_, and *s*_1_ respectively. The pink arrow shows immune stimulation by uninfected cancer cells, while the red arrow shows stimulation by infected cancer cells.

The values of the initial conditions are presented in Table 1.

**Table 1:** Initial conditions for model variables

| Variable | Value | Unit | Description |
| --- | --- | --- | --- |
| $C_0$ | $10^6$ | cells/mm<sup>3</sup> | Initial density of uninfected cells |
| $I_0$ | 0 | cells/mm<sup>3</sup> | Initial density of infected cells |
| $V_0$ | $1.9 \times 10^{10}$ | viruses/mm<sup>3</sup> | Initial density of virus particles |
| $Y_0$ | $10^4$ | units/mm<sup>3</sup> | Initial density of immune cells |

Pooladvand et al. [36] considered the carrying capacity of a solid tumor of radius 1 mm is about *L* = 10^6^ cells per mm^3^, this value is based on Lodish [29]. We consider this value as the initial density of uninfected tumor cells, *N*_0_, in our model. Based on the experiments by Kim et al. [24] on adenovirus in the glioblastoma U343 cell line, the tumor growth rate was estimated to be approximately 0.3 per day. This value was also adopted by Baabdullah et al. [4] in their model, and we will, likewise, use it in our analysis. The initial condition for the amount of adenovirus virions is *V*_0_ = 1.9 *×* 10^10^ virions per mm^3^. We also adopt the same parameter values for *L, β, ω*, and *b* as those used in the model by Baabdullah et al. [4]. These parameters are associated with a reovirus and were applied in the context of breast cancer. For the remaining parameters, we use estimates based on the immune-virotherapy interaction model developed by Al-Tuwairqi et al [2]. In their study, these parameters were examined in the context of glioma treatment. The virus-related parameters in their model were derived from the study by Friedman et al. [15], which focused on the mutant herpes simplex virus 1 (hrR3). Al-Tuwairqi’s immune model [2] incorporated components of the innate immune response, including natural killer cells and cytokine activity. We summarize the model parameters in Table 2. To simplify our analysis, we apply the non-dimensionalization method by considering

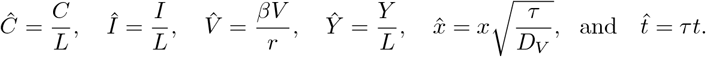

**Table 2:** Parameters for the base model (2), their baseline values, and their references.

| Parameter | Description | Baseline Value | Units | Source |
| --- | --- | --- | --- | --- |
| $r$ | Growth rate of uninfected tumor | 0.3 | (day)<sup>-1</sup> | [4] |
| $L$ | Tumor carrying capacity | $10^6(10^6 - 10^8)$ | cells | [4] |
| $\beta$ | Infection rate of uninfected tumor cells by OV | $1.5 \times 10^{-9}$ | (PFU .day)<sup>-1</sup> | [4] |
| $p_1$ | Killing rate of tumor by immune cells | $0.48 \times 10^{-6}$ | (cell .day)<sup>-1</sup> | [2](estimated) |
| $\alpha$ | Death rate of infected tumor cells | 1 | (day)<sup>-1</sup> | [4] |
| $q_1$ | Killing rate of infected tumor by tumor-specific immune cells | $0.63 \times 10^{-6}$ | (cell .day)<sup>-1</sup> | [2](estimated) |
| $b$ | Viruses released by lysed infected tumor cell | 3500 | (PFU)(cell)<sup>-1</sup> | [4] |
| $\omega$ | Virus-induced immune response rate | 4 | (day)<sup>-1</sup> | [4] |
| $s_1$ | Killing rate of virus by immune cells | $0.21 \times 10^{-6}$ | (cell .day)<sup>-1</sup> | [2](estimated) |
| $\eta_1$ | Stimulation of immune response by uninfected cells | $3.84 \times 10^{-7}$ | (cell .day)<sup>-1</sup> | [2](estimated) |
| $\zeta_1$ | Stimulation of immune response by infected cells | $7.8 \times 10^{-7}$ | (cell .day)<sup>-1</sup> | [2](estimated) |
| $\xi_1$ | clearance rate of immune cells | 0.036 | (day)<sup>-1</sup> | [2] |

The dimensional model (2) transforms into the non-dimensional system:

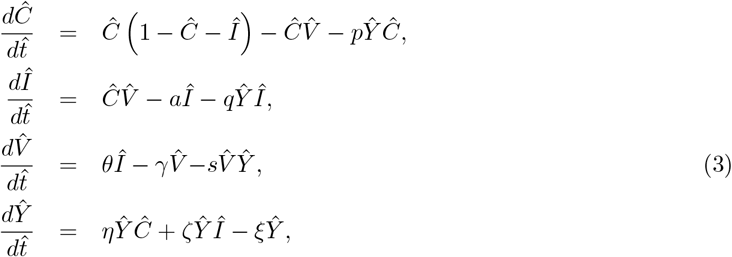

where

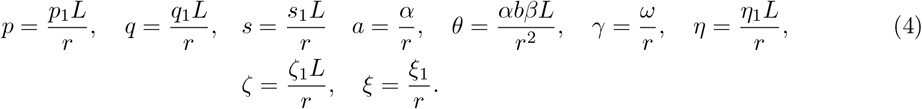

The parameter values before and after non-dimentionalization are given in Table (2) and (3), respectively.

Note that we solve (3) numerically and rescale the results into dimensional variables such that we present densities and time in their physical units.

## 3 Analysis of the Base Model

Here, we summarize the basic dynamical properties of Model (3). A complete analysis of the model is presented in [33]. Model (3) has the following equilibria:

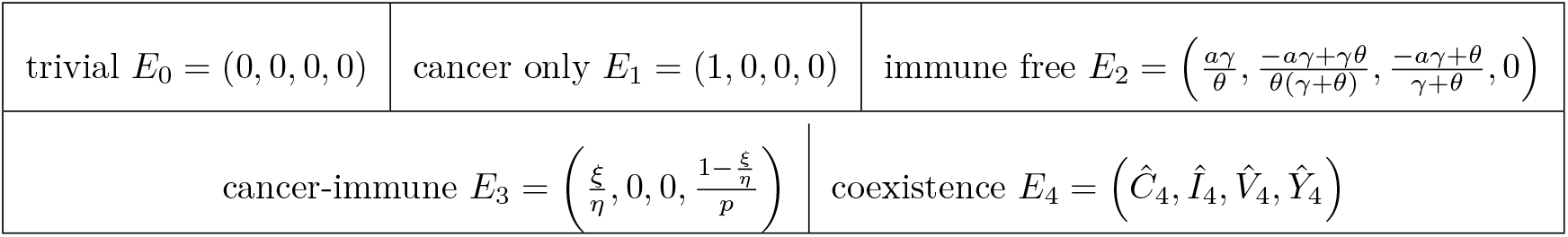

where *E*_4_ is the coexistence point which cannot be expressed explicitly. In [33], we prove the following Theorem.

### Theorem 1.

*For our base immune-oncolytic virus model* (3), *we have the following results:*

- *The basic reproduction number is given by* 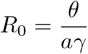,
- *E*_0_ *is always a saddle, and when the basic reproduction number R*_0_ *is less than one, or equivalently θ < aγ, E*_1_ *is locally asymptotically stable and E*_2_ *is not biologically relevant*,
- *When R*_0_ *is greater than one, E*_1_ *becomes unstable and the coexistence steady state arises through a transcritical bifurcation at θ*_*t*_ = *aγ, where E*_2_ *becomes biologically relevant*,
- *E*_2_ *is locally asymptotically stable when*

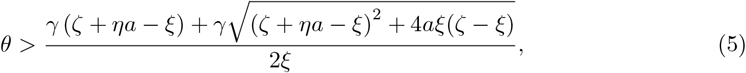

*and*

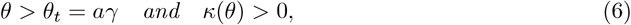

*where*

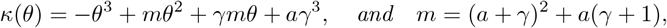
- *There exists a bifurcation value θ*_*H*_ *> θ*_*t*_ *with κ*(*θ*_*H*_) = 0 *such that the system undergoes a Hopf bifurcation at E*_2_,
- *E*_3_ *is locally asymptotically stable only if we have*

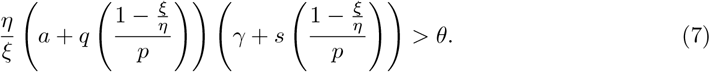

### 3.1 Tumour control probability

To effectively compare treatment strategies and better estimate their expected success, we need a suitable metric. Traditionally, tumour control probability (TCP) has been used in radiotherapy as a measure of the likelihood that treatment has successfully eliminated all clonogenic cells. In the study by Gong et al. [18], the Poisson-based tumor control probability (TCP) was identified as a reasonable first-order approximation for comparing the effectiveness of different treatment protocols. The Poissonian TCP is based on a stochastic process for the random variable *X*_*t*_ of viable cancer cells at time *t*. Assuming that individual cell deaths occur independently and that survival events are rare, *X*_*t*_ can be approximated by a Poisson distribution. We have that under a Poisson distribution with the expectation of *λ* events in a given interval, the probability of k tumour cells surviving is given by [47]

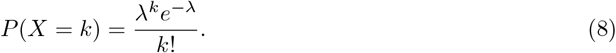

If *N*_0_ denotes the initial number of tumor cells and *S*(*t*) the surviving fraction at time *t*, then the expected number of surviving cells is given by *E*(*X*_*t*_) = *λ*. Approximating this expectation by *N*_0_*S*(*t*), we obtain *λ* = *N*_0_*S*(*t*). Consequently, the Poisson-based tumor control probability (TCP) is

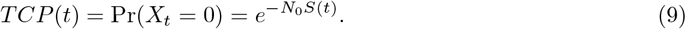

In our model (2), we normalize the cell density *C*(*t*) with *L* = *N*_0_. Hence, here the survival fraction is simply 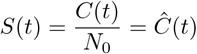. Therefore, our TCP is given by

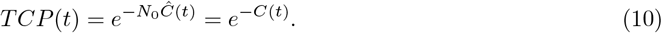

We use our base value parameters from Table (3) to numerically solve system (3), and compute the cell densities, virus load, and TCP in the first row of Figure 2.

**Table 3:**
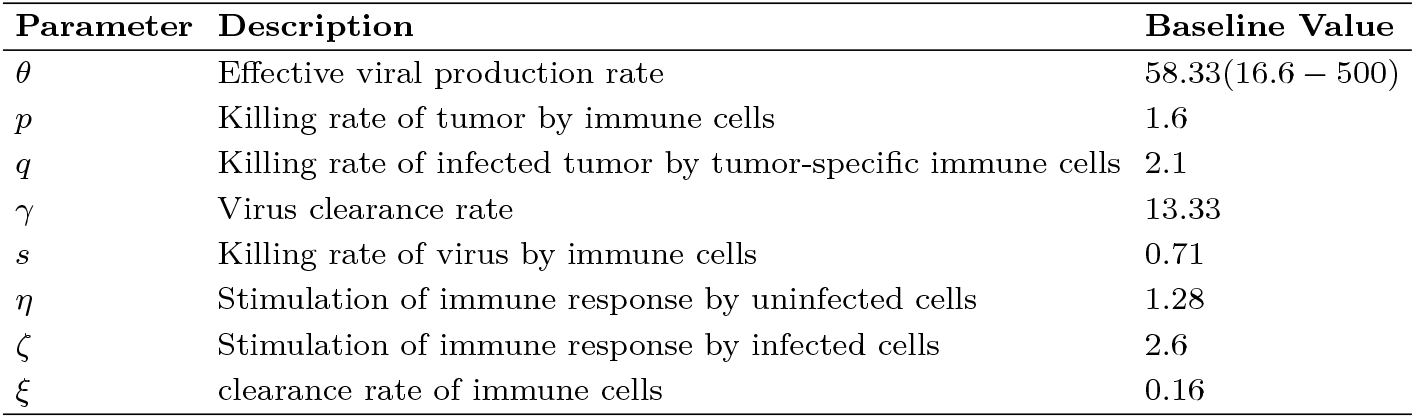
Parameters for the base model (2) and their baseline values after non-dimensianlization (4).

| Parameter | Description | Baseline Value |
| --- | --- | --- |
| $\theta$ | Effective viral production rate | 58.33(16.6 – 500) |
| $p$ | Killing rate of tumor by immune cells | 1.6 |
| $q$ | Killing rate of infected tumor by tumor-specific immune cells | 2.1 |
| $\gamma$ | Virus clearance rate | 13.33 |
| $s$ | Killing rate of virus by immune cells | 0.71 |
| $\eta$ | Stimulation of immune response by uninfected cells | 1.28 |
| $\zeta$ | Stimulation of immune response by infected cells | 2.6 |
| $\xi$ | clearance rate of immune cells | 0.16 |

**Figure 2:**
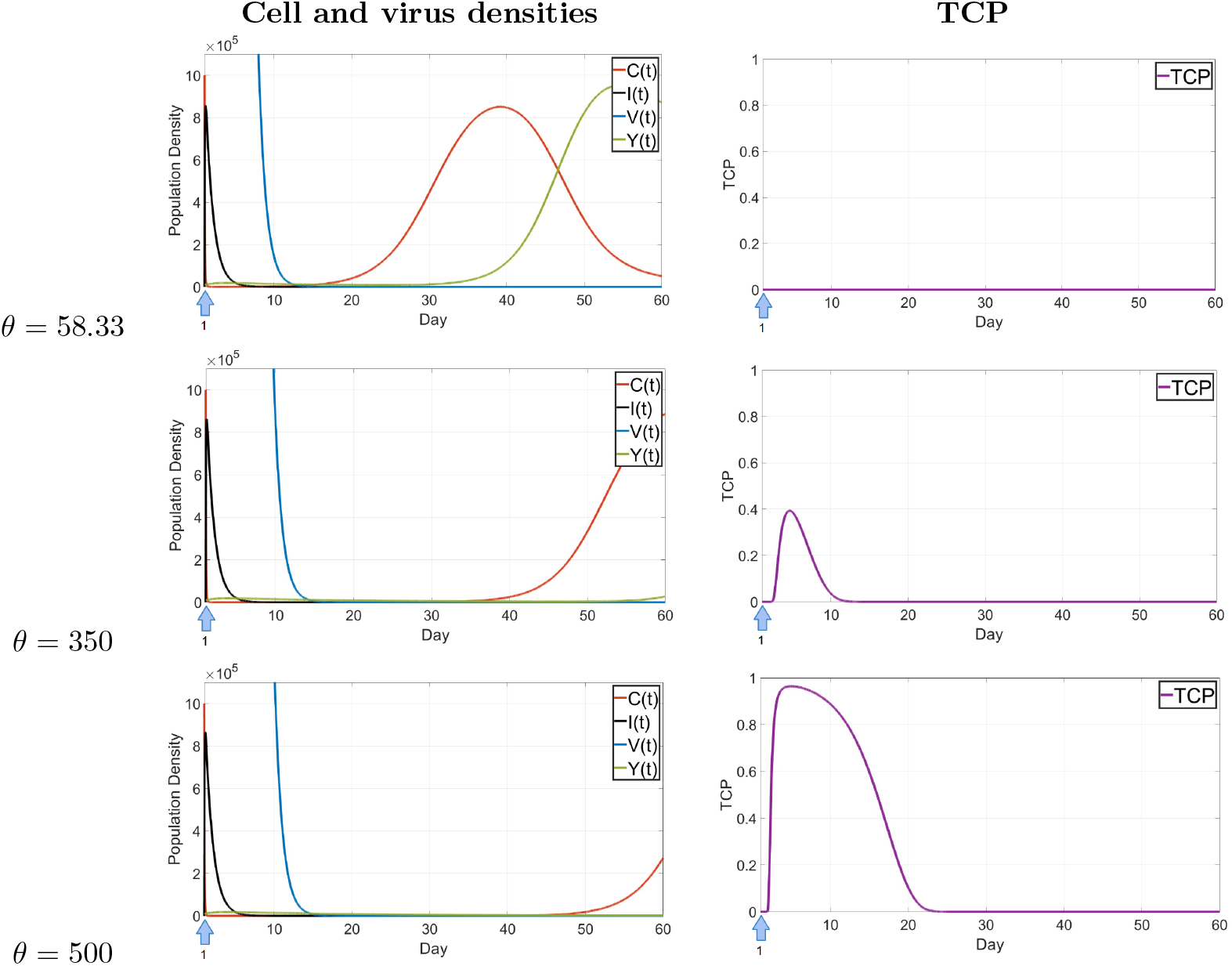
Strategy (S1): Dynamics of the cancer cells, virus, and immune cells of model (3) for a single dose of virus with different values of *θ*. The density of virus particles is re-scaled by dividing by 100. Note that we solve (3) numerically and rescale the results into dimensional variables so we can present densities and time in their physical units. The right column shows the corresponding TCP plots. The small blue arrow indicates the day of administration of the viral dose.

In our simulations, we also introduce a cut-off value to regulate the virus load. During simulations, if at any time the virus load drops below 10^*−*14^, it is considered cleared and is set to be zero. Otherwise the numerical solver would sometimes assign a negative value to *V*, and then the simulation breaks down.

The plots of the base case in the first row of Figure 2 indicate a rapid initial decline in cancer cells when the oncolytic virus (OV) is administered; however, the cancer cells quickly recover and approach their carrying capacity. Activated immune cells manage to reduce the population of cancer cells, but the tumor is not cleared. There is no signal in the TCP value throughout the base case, indicating that with these parameter values the OVT alone is not successful.

## 4 Model Strategies to Improve Treatment Efficacy

We observed in the previous section, that the mathematical model predicts OVT as a monotherapy with one viral injection would not be successful, confirming the clinically observed outcomes. In this section, our goal is to use the model to discuss improved treatment strategies. We will refine our model, explore different parameter values, and test various combination therapies to achieve a more effective approach for cancer eradication. One of our key strategies will be combining virotherapy with immunotherapy, as this has been recognized as one of the most promising ways to enhance OVT efficacy. At the end of this section, we will compare all the results to gain a better understanding of how virotherapy can be optimized for more successful treatment outcomes. Our base model will be system (3), where *Y* is considered a general immune response which acts on both the virus and cancer cells. We will build upon this model by incorporating enhancements to improve treatment effectiveness. Here is a quick look at the strategies we will consider in the following:

(S1) Increasing the effective viral production rate *θ*. The parameter *θ* is a combined quantity (see (4)), which includes the viral binding rate *β* and the burst size *b*. Increasing either of these parameters increases *θ*.
(S2) Developing optimized daily and weekly dosing schedules for up to four weeks following the first viral injection.
(S3) Combining virotherapy with immunotherapy:
  a. Immunosuppression strategies
  b. Immune stimulation approaches
  c. Combination with CAR-T cell therapy, including optimized scheduling strategies

### 4.1 Strategy (S1): Increase of viral production rate

The efficacy of the virus in eradicating cancer cells largely depends on its inherent properties, including the infection rate *β*, and burst size *b*. Therefore, we will closely examine the parameter 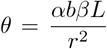 which incorporates both of these factors. In the estimations performed by [39], the burst size *b* of adenoviruses was found to range between 1, 000 *−* 100, 000. In our simulations shown in Figure 2, we use the baseline parameter values from Table (2), where *β* was considered to be 3500, and thus, *θ* was set to 58.33. According to the findings in [39], the possible range for *θ* extends to over 1600. In this section, we investigate whether increasing the effective production rate has a positive impact on the outcome of OVT.

As it is seen in Figure 2, increasing *θ* value in model (3) is an effective approach to enhance the efficacy of OVT. The therapy becomes relatively successful, with the TCP curve approaching one, only when *θ* is nearly ten times higher than our baseline value. While this strategy significantly improves OVT’s effectiveness and brings it closer to achieving complete cancer eradication, it is important to note that this value is currently out of reach for realistic viruses.

### 4.2 Strategy (S2): OVT schedule optimization

Related to the under performance of OVT, Ornella et al. [35] argued that lower doses distributed over several days, rather than the maximum tolerated dose, may be associated with improved tumor response. In [44], Naumenko et al. pointed out that despite a lack of direct infection, subsequent viral doses—through a monocyte-dependent mechanism—enhance and sustain the infection initiated by the first dose. This process enhances CD8^+^ T cell recruitment, delays tumor progression, and improves survival in multi-dosing OV therapy. These findings highlight the importance of dose optimization for the successful clinical development of OVT. Having a mathematical model available (3) we now address such a scheduling optimization. Our objective is to identify treatment schedules that minimize tumor load at the end of therapy, improving upon the results observed previously. The goal is to increase TCP values and maintain them at 1 for an extended period.

We incorporate a control term *u*(*t*) into our model (3) to analyze treatment success. We test various dosing schedule with different viral dose levels. Our analysis considers treatment durations ranging from one week to five doses over a four-week period. A key constraint in our optimization is that the total dose administered each week must be equal to 95, which is equal to the initial value that was used by Baabdullah et al. [4]. Adding a control term *u*(*t*) in the non-dimensional model (3) gives

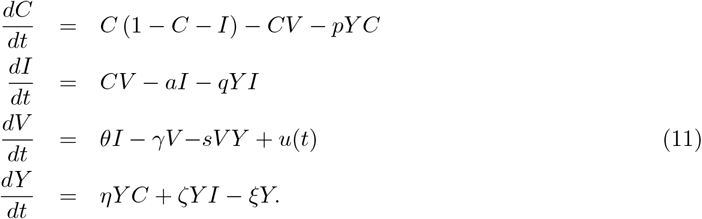

We focus on treatment schedules that can be realistically implemented in clinical practice. Specifically, we allow at most one treatment per day and exclude treatments on weekends. We set the treatment to be performed in a period of 4 weeks or 28 days. However, it is important to note that the model (11) is nondimensionalized, and the nondimensional time 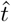 is defined by 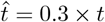. This implies that day 28 corresponds to 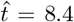. Consequently, the objective function of this optimization problem is to minimize the uninfected tumor load 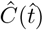 at the end of the period 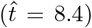. For simplicity, we will omit the hats from this point forward, with the understanding that all variables refer to their nondimensionalized form unless otherwise specified. We let *N* denote the total number of doses and *S* represent the schedule matrix. *S* has 4 rows and 5 columns. Each row stands for a week of therapy and each column corresponds to a day of the week. Since no therapy is performed on the weekends, we only consider 5 columns– Monday, Tuesday, Wednesday, Thursday, and Friday. The matrix *S* consists of *0’s* and *1’s*. We set an element of the matrix to be 1 if a dose of virus is administered on the corresponding day noting that we do not consider multiple doses on a single day. We also consider a matrix *D* with the same size as *S*, representing the value of dose given on the corresponding day. The *schedule matrix S* and the *dosage matrix D* can be written as

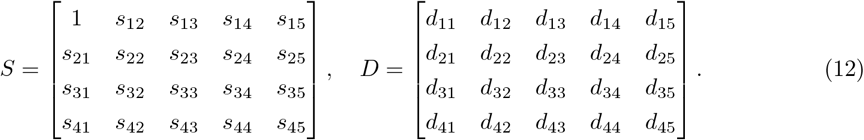

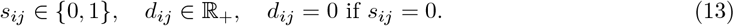

The element (1, 1) is considered to be 1, since we assume we always perform virotherapy on day 1, which in this version correspond to *t* = 0.3. We define the set of suitable treatment schedules *U*, as the set of all the *u*(*t*) belonging to the measure space *M* [21] defined as

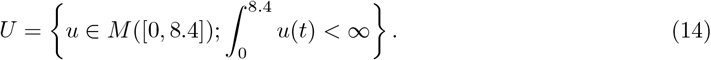

We also add the constraint that the total dose on each week should be equal to 95 and we write it as

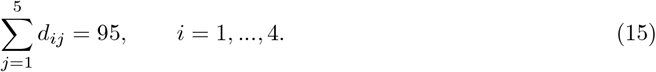

We define our specific treatment schedule *u*(*t*) as

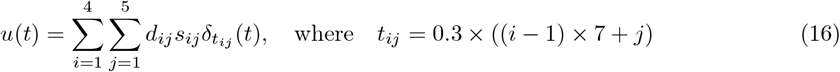

and 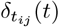 denotes the Dirac-delta distribution and *u*(*t*) *∈ U* belongs to a set of suitable treatments as defined in (14). The *t*_*ij*_ represents the days the treatment is administered starting from 1 to 28. In our optimization problem, we optimize the matrix *D* and *S* such that at the end of the period, *C*(*t* = 8.4) is minimized.

#### Example 2.

*Consider a treatment schedule such that the virotherapy is applied on the Monday of week one with dose 95, and Thursday and Friday of week three, and Monday and Tuesday of week four, each with dose 47*.*5. The suitable values of each dose are obtained through the optimization simulation and thus the corresponding schedule and dosage matrices S and D are written as*

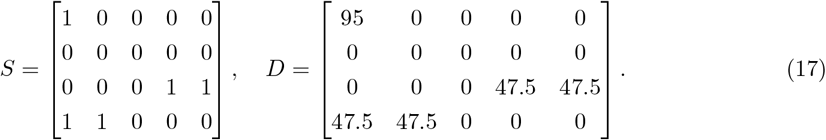

For the optimization we fix a schedule matrix *S* and we optimize for the dosage matrix *D*. We find a large number of good schedules and we show some examples in Figures 3 and 4.

**Figure 3:**
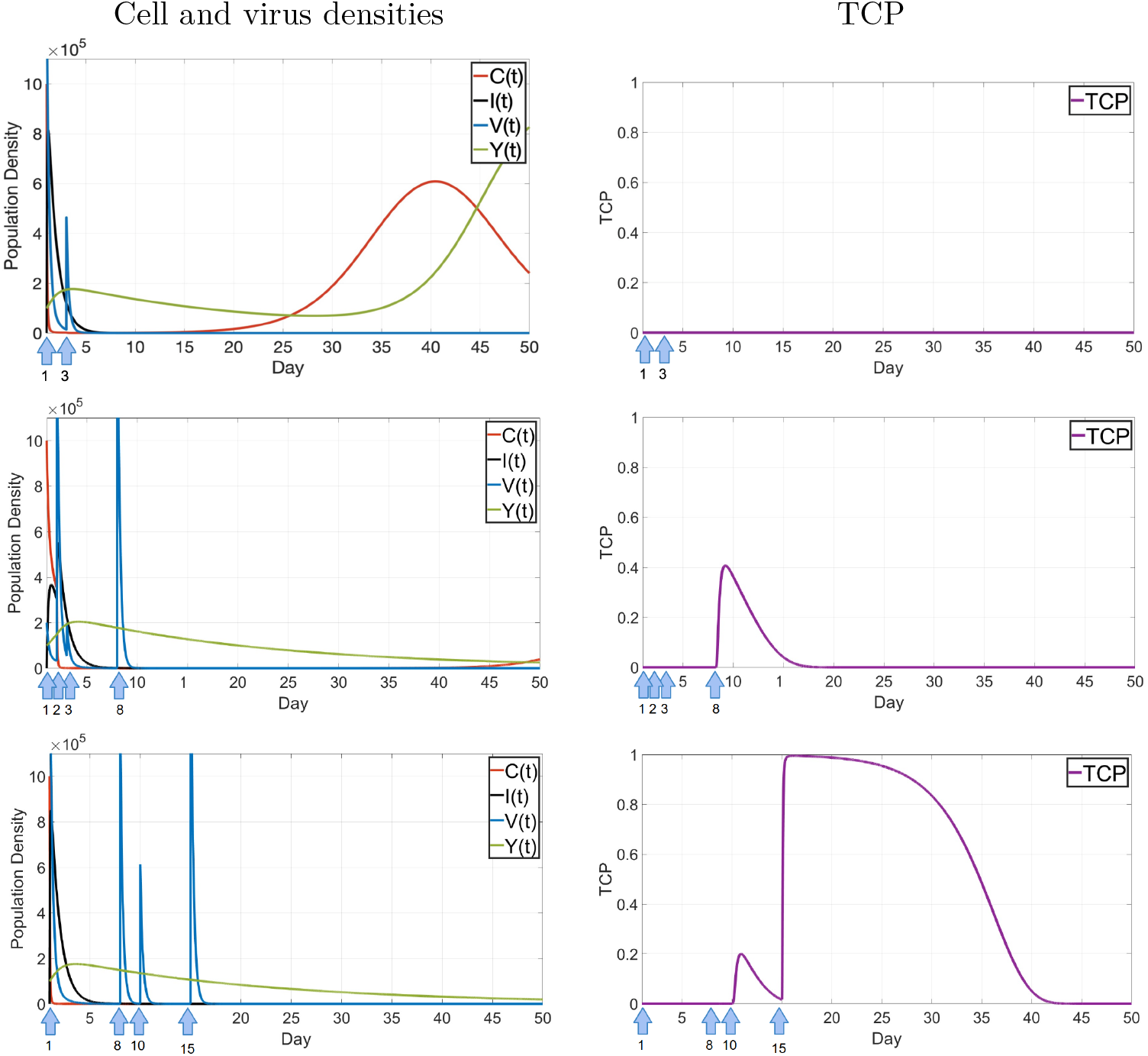
Strategy (S2), optimized scheduling. Examples of some of the optimized treatments. Left shows the dynamics of the cancer cells, virus, and immune cells and right the corresponding TCP. The density of virus particles are re-scaled by dividing by 100. The first row shows treatments on days 1 (with dose 72.50) and 3 (22.50), second row on days 1 (10), 2 (77.76), 3 (7.24), and 8 (95), and third row days 1 (95), 8(64.35), 10 (30.65), 15 (95). Note that we solve (3) numerically and rescale the results into dimensional variables such that we present densities and time in their physical units.

**Figure 4:**
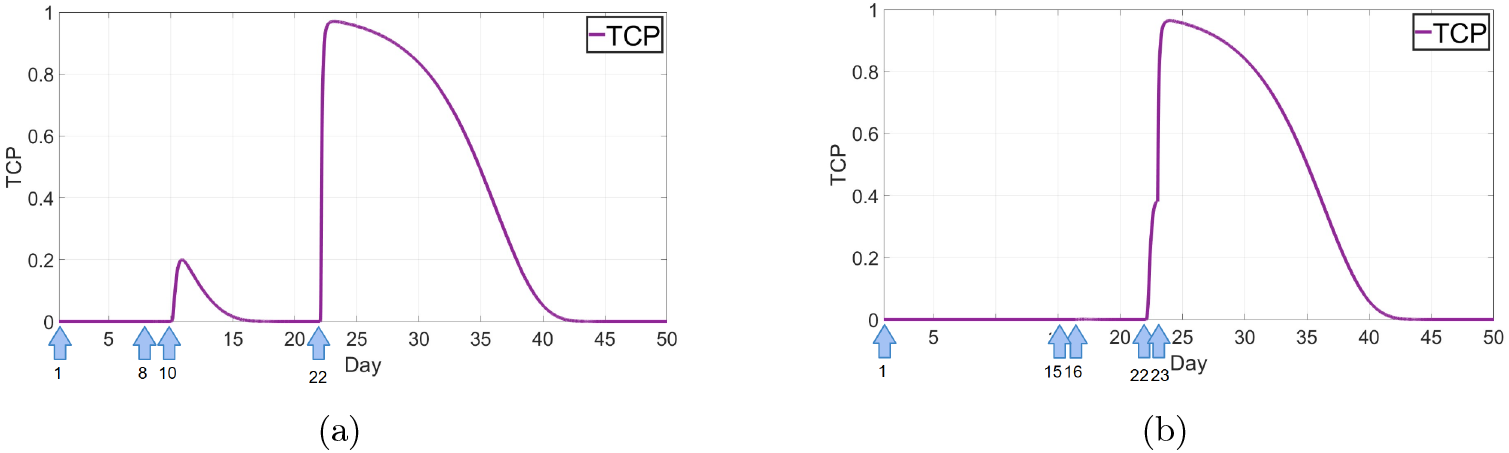
Strategy (S2), optimized scheduling. (a): The TCP curve of 4 doses of virus of value of 285 on days 1(value 95) day 8(value 64.86), day 10(value 3.14), and day 22 (value 95). (b): The TCP curve of 5 doses of virus of value of 285 on days 1(value 95) day 15(value 47.50), day 16(value 47.50), day 22 (value 47.47), and day 23 (value 47.53).

In the first case in Figure 3, we distribute the initial viral dose of 95 over the first week of therapy. The cancer cell count remains low for some time period during the first week, which is slightly longer than the case of treatment on day 1 (Figure 2 top). While this schedule shows partial improvement, the cancer is not fully eradicated, and as a result, the TCP value remains at zero throughout the entire period. Next, in the middle row of Figure 3, we consider treatments in week one and two. A total dose of 190 is administered over two weeks, with four doses in total (95 per week). After the fourth dose on day 8, the cancer curve remains at zero for almost 30 days and the TCP shows a signal. A clear improvement compared to Figure 2 top.

To further investigate the effects of prolonged treatment, we distribute a total of 285 doses over three weeks in Figure 3 and over four weeks in Figure 4. In the three-week case, we see a clear treatment success after day 15, where the TCP raises to 1. In the four-week treatments, we see good outcomes after 22 days.

We optimize many more schedules and we find a large number of successful schedules. We summarize them as a heat map in Figure 5. Each line in Figure 5 presents a schedule, where the days per week are the columns. On a treatment day, we use a color from dark red for a low dose to yellow for a high dose. The last column indicates the resulting maximum TCP. Bright yellow indicates successful cases. This figure illustrate the impact of various dosing schedules on the tumor control probability, highlighting how prolonged and optimized administration influences treatment outcomes. This aligns with the findings of Naumenko et al. [44], who suggested that while subsequent treatments may not have a direct infection effect, they enhance and sustain the infection initiated by the first viral dose, promote immune cell recruitment, and delay tumor growth.

**Figure 5:**
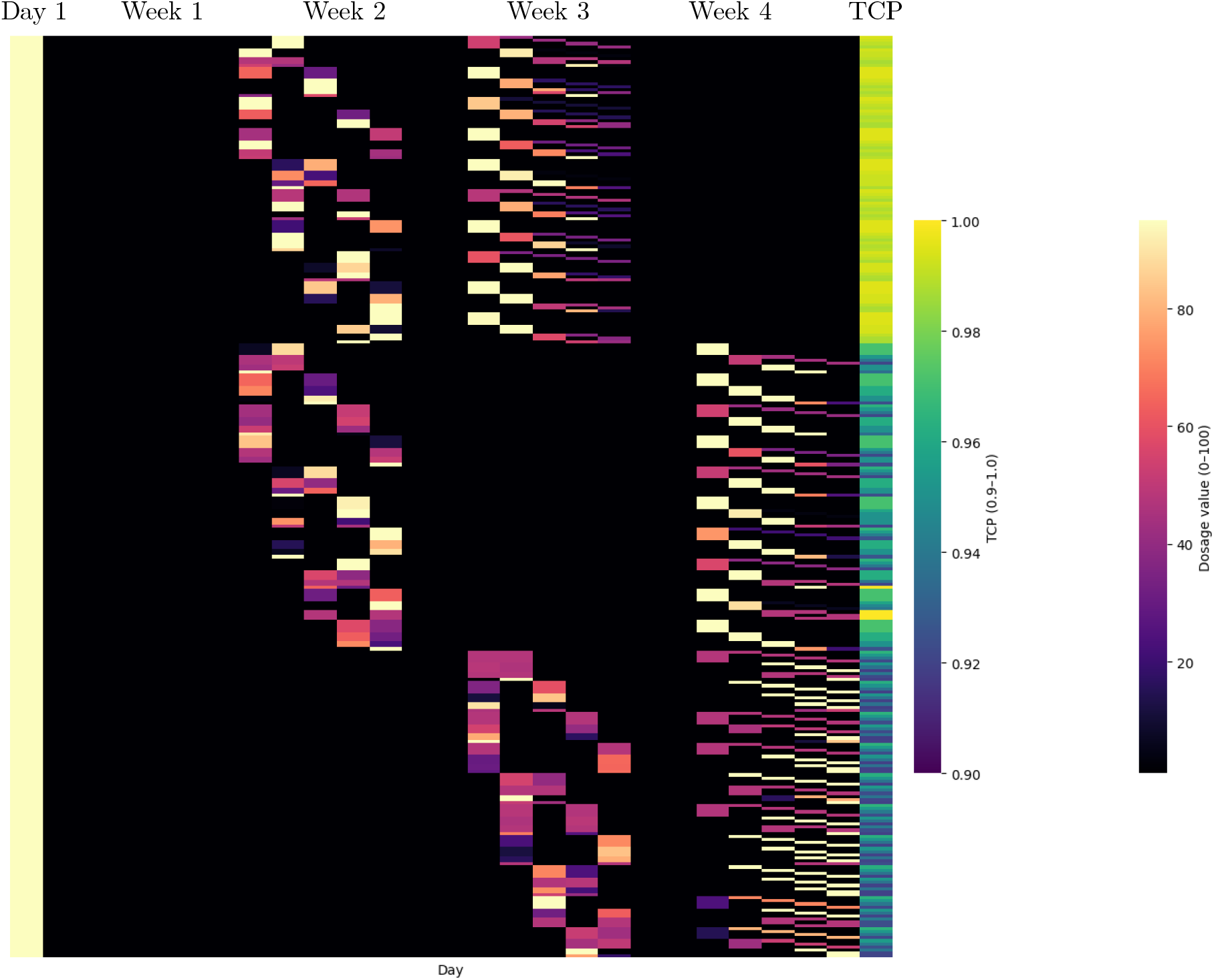
Summary of treatment schedules alongside their corresponding optimized dose values and the highest TCP values over a four-week period. Each line in represents a schedule, where the days per week are the columns. On a treatment day we use a color from dark red for a low dose to yellow for a high dose. The last column indicates the resulting maximum TCP.

### 4.3 Strategies (S3): Combination of virotherapy and immunotherapies

As a result of tumor cell infection with OVs, an antitumor immune response is activated, transforming immunologically “cold” tumors into “hot” ones [16]. The lysis of tumor cells induced by OVs releases tumor antigenes and stimulates the immune system, triggering immunogenic cell death. This process leads to the release of danger-associated molecular patterns (DAMPs and PAMPs) and innate immunity signaling molecules such as HMGB1 and ADP. Consequently, the immunosuppressive tumor microenvironment (TME) shifts to a proinflammatory state, activating innate immunity in nearby cells [7]. A combination of OVT and immuno therapies opens new avenues for developing more effective treatment regimens [1]. In this section, we will examine the efficacy of combining oncolytic virotherapy (OVT) with immune suppression, immune stimulation and with chimeric antigen receptor (CAR) T-cell therapy.

#### 4.3.1 Strategy (S3a) Immunosuppression

The activation of the immune system may significantly limit viral replication and spread. This ultimately causes the reduction of the therapeutic efficacy of OVs [3]. The use of immunosuppressive drugs could be a strategy to inhibit this effect. There has also been studies, which have shown that immunosuppressive treatment at the time of OVT did confer a negative impact on cancer survival and there was a trend towards worse overall survival, although not statistically significant [43]. In our model (3) we applied the immunosuppressive effect of the drugs with the factor *ϵ*_*t*_. Our modified model is given as follows

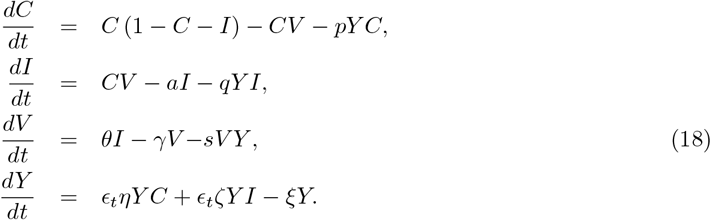

We consider the effect of the drug only on the stimulation terms and not on the clearance rate. We study the results of the immunosuppression treatment combined with oncolytic virotherapy for different values of *ϵ*_*t*_ in Figure 6, and we do not observe any improvement in the TCP values of the system. We observe that the TCP values stayed zero (plots not shown) and thus the combination of these two treatments is one of the strategies that would not improve the outcome of OVT, according to our model.

**Figure 6:**
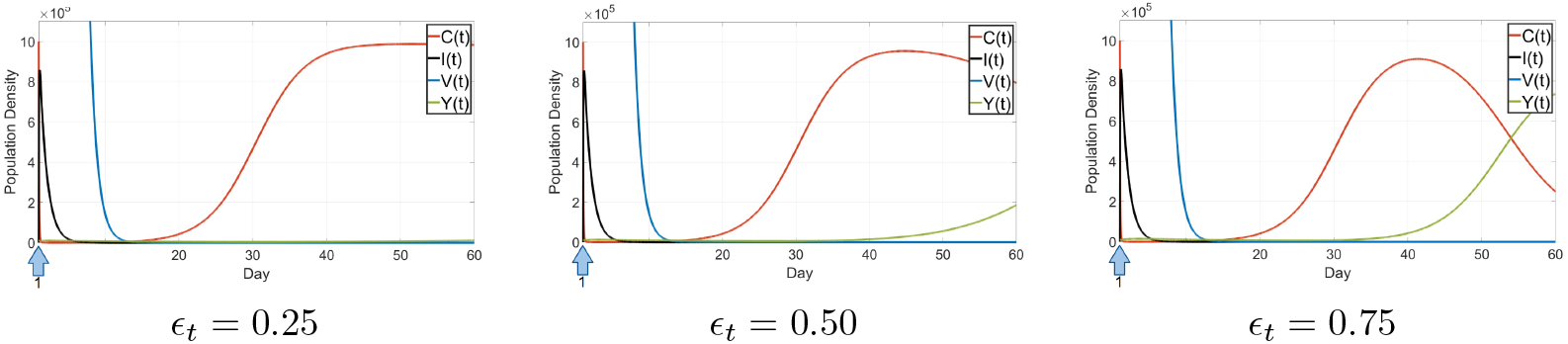
Strategy (S3a) Immunosuppression: Dynamics of the infected and uninfected cancer cells, the virus, and the immune cells of model (18) in combination therapy of immunosuppressive drugs and oncolytic virotherapy for different values of *ϵ*_*t*_. Again, the solutions are presented in dimensional coordinates.

#### 4.3.2 Strategy (S3b): immune stimulation

One of the promising strategies of oncolytic virotherapy (OVs) enhancement is to combine them with the use of T-cell engager molecules. This stimulates existing T cells to lyse tumor cells [17]. Scientists might call the immune modulators as the gas pedals of the immune system. Different types of therapies have been developed that improve the immune system’s ability to attack and eliminate cancer [8].

Similarly we add the factor 𝒳 of immune stimulator to the model (3). The model is given in equation (19).

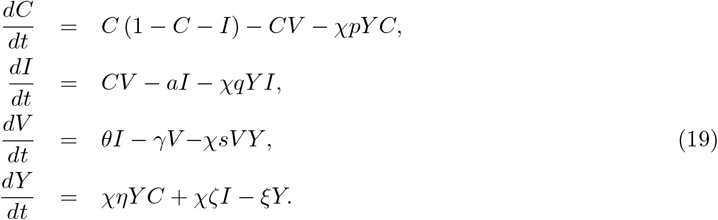

The results we obtained showed no improvement in the outcome of the treatment. We see in Figure (7), after modulating the immune system by a factor of 2.5, we observe that it temporarily reduces the tumor load in an oscillatory manner; however, the TCP plot remains sustained at zero (plots not shown).While a stronger immune response may more effectively target tumor cells, it can also increase susceptibility to pathogens. Numerous studies have reported that following OVT, significant amounts of the virus often accumulate in the spleen and liver [17]. An even more robust immune system would likely amplify this effect further.

**Figure 7:**
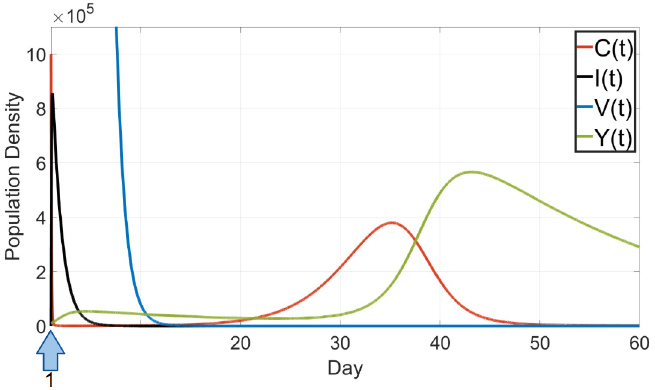
Strategy (S3b) immune stimulation: Dynamics of the infected and uninfected cancer cells, the virus, and the immune cells in combination therapy of immunostimulation drugs and oncolytic virotherapy for model (19).

#### 4.3.3 Strategy (S3c): Combination with CAR-T cells therapy

Our next strategy for improving the OVT efficacy is its combination with CAR-T cell therapy. In this therapy, in order to help T-cells recognize tumour-derived antigens, T-cells are extracted from the patient’s body, genetically engineered to express chimeric antigens receptors (CARs) and then they are infused back to the patient [6]. However, it has been shown that tumour-mediated immunosuppression is one of the biggest challenges in CAR-T cell therapy [40]. By combining OVT and CAR-T cell therapy, oncolytic viruses infect the tumour cells and make infected cells express target tumour antigens that is recognized by CAR-T cells [26].

Figure 8 helps to have a better understanding of how the infected, uninfected cancer cells interact with the virus and the CAR-T cells. Our base CAR-T cell ODE model in dimensional form is as follows

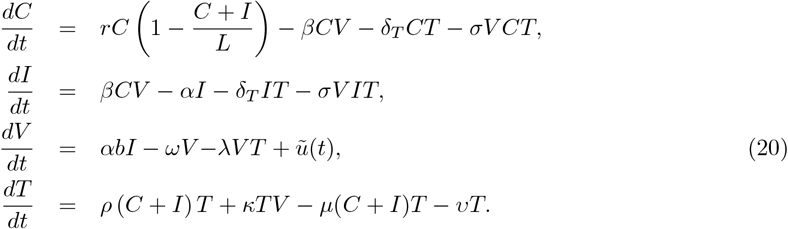

**Figure 8:**
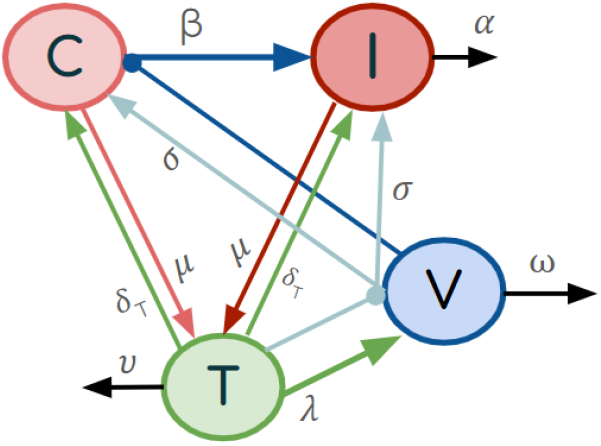
Sketch of the OVT-CAR-T model. The blue arrows represent the viral infection of cancer cells at a rate of *β*. The light cyan arrows indicate the virus-induced cancer cell killing of the *T* cells at rate *σ*, while the orange arrows illustrates their direct killing of the virus and the cancer cells at rates *λ* and *δ*_*T*_ respectively. The pink and red arrows denote denote the CAR-T cell death by the cancer cells at rate *µ*. All black arrows depict cell death and virus degradation.

In this model, similar to our base model (2), we consider *C* and *I* to represent the populations of uninfected and infected cancer cells, respectively. We introduce *T* as both the CAR-T cell population and the general immune cells that targets both cancer cells and free viruses— noting that effector CAR-T cells alone, only target cancer cells. Additionally, the population of free viruses (*V*) is represented in the third equation. The first two terms in equation one are similar as in model (2). The third and forth terms account for tumor cell killing by the effector CAR-T cells at rates *δ*_*T*_ and *σ* respectively, but the difference is that the fourth term denotes the OV-induced CAR-T killing. Terms one and two in the infected cancer cells equation are the same as in equation (2) and the two following terms represent the CAR-T cell killing similar to the first equation. In equation three, we have the effective replication term of the virus and their clearance in terms one and two respectively and the last term denotes the killing of the OV by the immune cells. *ũ*(*t*) is the dimensional control term and has a similar definition as (16) after non-dimensionalization. *ρ* represents the rate at which CAR-T/immune cells are stimulated and proliferate upon interacting with both infected and uninfected cancer cells, as described in the first term of equation four. The second term represents an OV-induced recruitment of CAR-T cells at rate *κ* inspired from [23]. The third term denotes the inactivation of CAR-T cells by both types of the tumor cells at the rate *µ* and their natural death at rate *υ* is accounted for in the last term.

In this model, the similar parameters as the equation (2) have the same values as in Table (2). The rest of parameter values are given in Table (4).

**Table 4:**
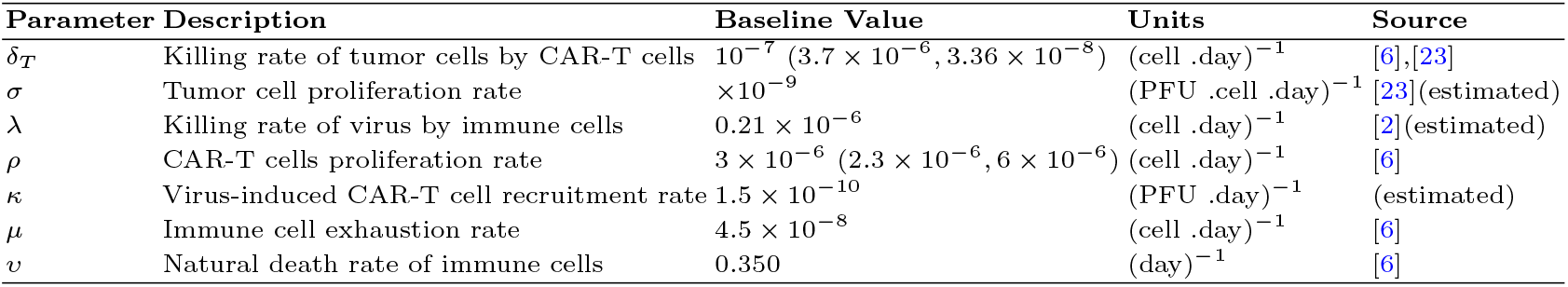
Model parameters for the onco-virus CAR-T cell model (20), their baseline values, units and references. Parameters not listed here are given in Table 2.

The parameters that are taken from Mahasa et al. [23] are from their study of glioma and the hrR3 virus. The CAR-T cell parameters from Barros et al. [6] are taken from their study on CAR-T cell therapy for mice with Hodgkin’s disease.

We set the initial conditions to be

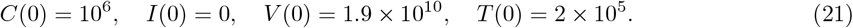

In Figure (9) we show several cases of treatment timing. In the first row of (9), we first simulate the behaviour of model (20) after non-dimenstionalization in the absence of virotherapy 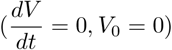.

**Figure 9:**
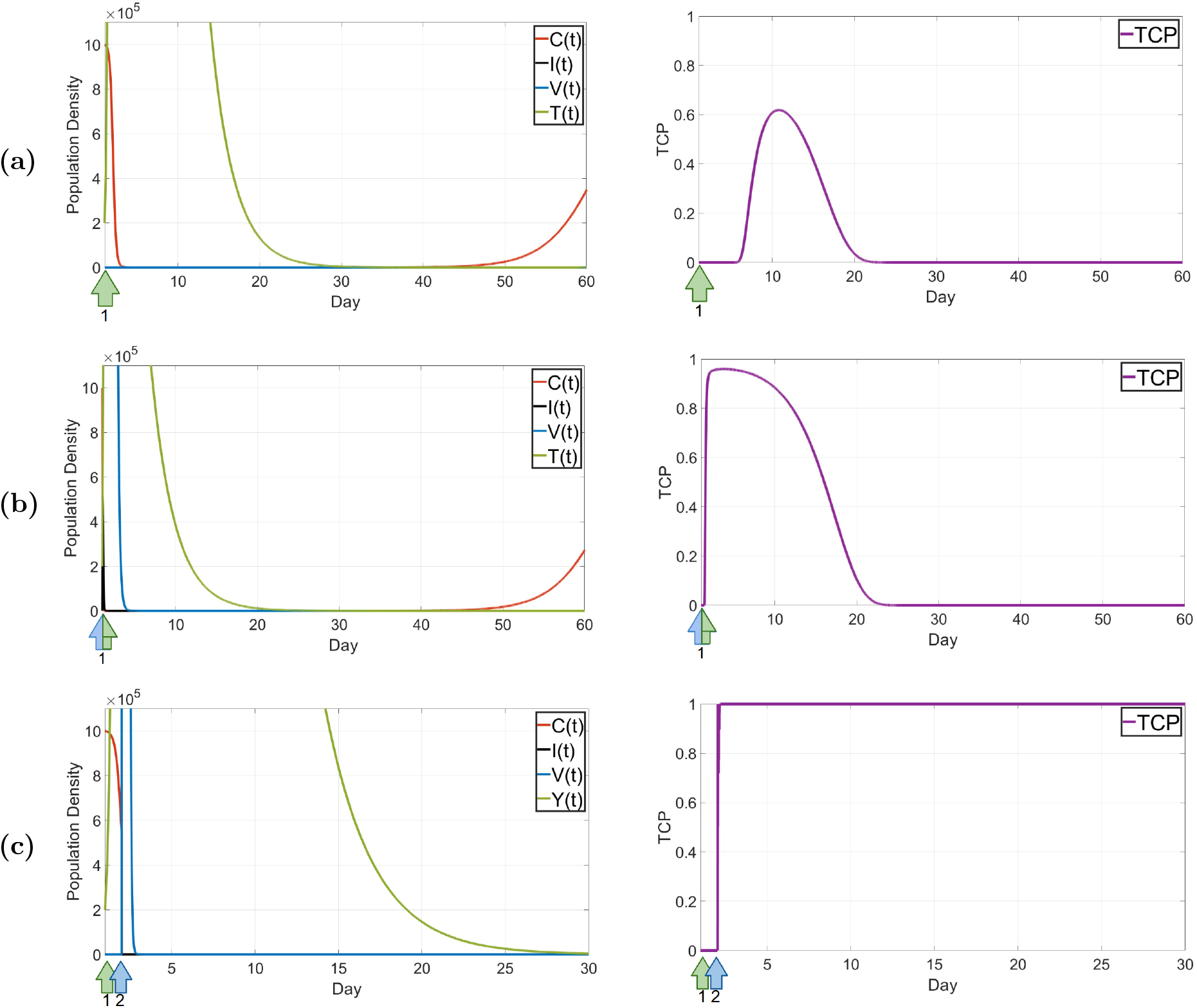
Strategy (S3c) combined with CAR-T: Dynamics of cancer cells and CAR-T cells in model (20) and the corresponding TCP. **(a)** A single dose of CAR-T cells without oncolytic virus. **(b)** CAR-T cells and oncolytic virus both administered on day one. **(c)** Optimized treatment: CAR-T therapy on day one and virus injection on day two.

We start the simulations without considering any optimization, and thus, the control term *ũ*(*t*) in (20) is zero. In Figure 9a, we see that although this monotherapy brings the cancer cells significantly down for a long time, the TCP curve only rises to slightly over 0.6. Combination of these two therapies is one of the suggested strategies to improve the efficacy of the treatments [13, 9, 20]. As oncolytic viruses (OVs) are armed with therapeutic transgenes, this causes a boost in the CAR-T activation. Also, in the immunosuppressed tumour’s microenvironment, OVs could be able to survive and even maintain their cytotoxicity functions [12]. When combined, we see in Figure 9b that the TCP value has a quick jump to about 0.98 just by an injection of CAR-T cells and the OV on day one. This shows the importance of the combination of virotherapy with immunotherapy for treatment efficacy and patients benefit.

We apply a similar optimizing strategy as in model (11) by considering similar restrictions for the combination of CAR-T cells and virotherpy model (20) after non-dimensionalization. The goal is to improve the results we obtained from the non-optimized combination therapy.

We test different schedules to obtain a more optimal boost for this combination therapy. During optimization simulations, we observe that by a slight delay in the administration of the initial virus dose, a more minimized tumor load is achieved as this delay significantly improved treatment outcomes. As we see in Figure 9 row (c), When CAR-T cell therapy is administered on day one, followed by oncolytic virus treatment on day two, the outcome is significantly more effective. The initial CAR-T cell injection reduces the cancer cell population. Once the tumor burden is lowered, virotherapy starts. At this stage, in addition to directly infecting cancer cells, oncolytic viruses (OVs) act as a booster for CAR-T cells as well.

We summarize the outcome of our various strategies in Table 5. Successful treatments are highlighted in teal blue.

**Table 5:** Summary of treatment strategies, schedules, viral doses, and TCP for model (2). The most promising combinations are highlighted in green.

| Strategy | Treatment Week(s) | Schedule (Days) | Total Dose | Max TCP |
| --- | --- | --- | --- | --- |
| (S1) Increasing $\theta$ | Week 1 | Day 1 ($\theta = 58$) | 95 | 0 |
| | Week 1 | Day 1 ($\theta = 350$) | 95 | 0.39 |
| | Week 1 | Day 1 ($\theta = 500$) | 95 | 0.96 |
| (S2) Optimized schedule | Week 1 | Days 1, 3, 5 | 95 | 0 |
|  | Weeks 1, 2 | Days 1, 2, 3, 8 | 190 | 0.40 |
|  | Weeks 1, 2, 3 | Days 1, 8, 10, 15 | 285 | 0.99 |
|  | Weeks 1, 2, 4 | Days 1, 15, 16, 22, 23 | 285 | 0.97 |
|  | Weeks 1, 3, 4 | Days 1, 8, 9, 22 | 285 | 0.96 |
| **Combination Therapies** |  |  |  |  |
| (S3a) Immunosuppression ($\delta$) | Week 1 | Day 1 ($\delta = 0.25$) | 95 | 0 |
| | Week 1 | Day 1 ($\delta = 0.5$) | 95 | 0 |
| | Week 1 | Day 1 ($\delta = 0.75$) | 95 | 0 |
| (S3b) Immunostimulation ($\chi = 2.5$) | Week 1 | Day 1 | 95 | 0 |
| (S3c) CAR-T Model (A) | Week 1 | CAR-T: Day 1 | CAR-T: 2 | 0 |
|  | Week 1 | CAR-T: Day 1, OV: Day 1 | CAR-T: 2, OV: 95 | 0.62 |
|  | Week 1 | CAR-T: Day 1, OV: Day 0.2 | CAR-T: 2, OV: 95 | 1 |
| (S3c) CAR-T Model (B) | Week 1 | CAR-T: Day 1 | CAR-T: 2 | 0 |
|  | Week 1 | CAR-T: Day 1, OV: Day 1 | CAR-T: 2, OV: 95 | 0 |
|  | Week 1 | CAR-T: Day 1, OV: Day 2 | CAR-T: 2, OV: 95 | 1 |

## 5 Model Extensions

In the above models, we described immune effects through one global immune compartment that comprises of immune cells that react to cancer cells, to viral antigens, and include CAR-T cells. However, the immune interaction is more specific, and in the model extensions discussed here, we consider specific immune responses according to their actions on cancer cells, on viral antigens and for CAR-T cells. We find essentially the same results as before, i.e. a careful choice of a viral treatment schedule has a huge impact on the treatment outcome. We summarize the results here, and we provide the details in Appendix A.

### 5.1 Separate compartments for CAR-T cells and general immune cells

In model (22) we extend the model presented in (20) by distinguishing between general immune cells and CAR-T cells. The resulting five-dimensional system is discussed in appendix A.1. The outcome closely resembles what we observe in Figure 11a: CAR-T cells initially reduce the cancer cell population rapidly, yet this effect is insufficient to raise the TCP curve above 0.6. We then proceed by combining virotherapy with CAR-T therapy. As shown in Figure 11b, the TCP remains at zero when the two therapies are administered simultaneously. This outcome arises because the rapid action of oncolytic virotherapy eliminates cancer cells too quickly, limiting the opportunity for stimulation of CAR-T cell proliferation. By implementing our optimization strategy, which introduces a short delay in the administration of OVT, we allow CAR-T cells sufficient time to become effectively stimulated. Consequently, the virotherapy not only acts directly against cancer cells but also amplifies the cytotoxic response of CAR-T cells. In Figure 11c, we observe that introducing just a one-day delay in virotherapy administration boosts the TCP value to 1.

### 5.2 Model with virus and cancer specific immune cells

Our next step is to extend the base model (2) by separating the immune response into cancer specific and viral specific. The equations are given in in the Appendix in A.2. Similar to the base model 2, we analyze the same strategies as used above (see Section 4). The results are essentially the same as before, and we summarize those in Table 6.

**Table 6:** Summary of treatment strategies, schedules, viral doses, and TCP outcomes for model (24). Entries highlighted in green indicate the most promising scenarios.

| Strategy | Treatment Week(s) | Treatment Schedule (Days) | Total Dose | Max TCP |
| --- | --- | --- | --- | --- |
| (S1) Increasing $\theta$ | Week 1 | Day 1 | 95 | 0 |
|  | Week 1 | Day 1 | 95 | 0.39 |
|  | Week 1 | Day 1 | 95 | 0.96 |
| (S2) Optimized OVT schedule | Week 1 | Days 1, 3, 5 | 95 | 0 |
|  | Weeks 1, 2 | Days 1, 5, 11 | 190 | 0.53 |
|  | Weeks 1–3 | Days 1, 8, 9, 15, 16 | 285 | 0.99 |
|  | Weeks 1, 2, 4 | Days 1, 8, 9, 22 | 285 | 0.97 |
|  | Weeks 1, 3, 4 | Days 1, 15, 16, 22, 23 | 285 | 0.96 |
| **Combination Therapies** |  |  |  |  |
| (S3a) Immunosuppression ($\delta$) | Week 1 | Day 1 | 95 | 0 |
|  | Week 1 | Day 1 | 95 | 0 |
|  | Week 1 | Day 1 | 95 | 0 |
| (S3b) Immunostimulation ($\chi = 2.5$) | Week 1 | Day 1 | 95 | 0 |
| (S3c) CAR-T therapies | Week 1 | CAR-T: Day 1 | CAR-T: 0.2 | 0 |
|  | Week 1 | CAR-T: Day 1; OV: Day 1 | CAR-T: 0.2; OV: 95 | 0.99 |

## 6 Conclusions

Using a virus to treat cancer is an intriguing idea. Research on oncolytic viruses is at full swing and some have made it already through clinical trials and FDA approvals [19]. However, the efficacy of OV therapy, even for approved viruses, is rather limited because of physical barriers, tumor heterogeneity, and an immunosuppressive tumour microenvironment [38]. Several strategies to improve the efficacy of OV therapy are currently considered. Researchers genetically modify viruses to make them stronger, more or less infective, diffuse faster, and reproduce better [5]. The combination of OV with different methods of immunotherapy has been extensively studied because OV is a natural activator of both innate and adaptive immunity [34]. However, treatment scheduling as an approach to increase the efficacy of OV therapy, has so far been ignored. Here, we use a mathematical model to compare OVT and OVT combination therapies with optimized treatment scheduling.

Inspired by the model proposed by Al-Tuwairqi et al. [2], we develop a model that captures the dynamics between cancer cells, oncolytic viruses, and the immune system. Our analysis confirmed earlier findings that oncolytic virotherapy (OVT), when used as a monotherapy at baseline values, is insufficient to fully eradicate cancer cells or prevent tumor relapse following treatment. To enhance the therapeutic outcome of OVT, we explored several strategies. We first examined the effect of increasing the effective viral production rate. Although this demonstrated a significant improvement in therapeutic outcomes, achieving such levels remains unrealistic with the viral strains currently existing in clinical use. A particularly novel contribution of this study was the implementation of optimized dosing and scheduling of OVT within our model. Simulation results highlighted the critical importance of secondary dosing. In fact we found a large set of effective treatment schedules (see Figure 5), each involving OV treatments that could even be two weeks apart from each other.

Notably, the therapeutic benefit of OVT following the initial dose was not primarily due to direct infection of target cells, but rather its role in stimulating and boosting the immune response. Our final strategy involved combining OVT with immunotherapies. The first combination tested included immunosuppression and immune stimulation. Neither approach enhanced the effectiveness of OVT: in the case of immunosuppression, OVT remained too weak to eradicate the infection on its own; in contrast, immune stimulation alone cleared the virus more aggressively, undermining the potential benefits of OV therapy. Our final combination therapy, was the combination of OVT with CAR-T cell therapy. Using our optimization framework, we found that the most effective treatment outcome occurred when OVT was applied a day after immune therapy were administered. This was because simultaneous application hindered proper stimulation of CAR-T cells by initial rapid reduction of cancer cells. A summary of these results is given in Table 5.

To investigate the robustness of our findings, we extended our model by explicitly distinguishing between general immune cells and also immune cells and CAR-T cells. This extended version produced results consistent with those of our base model, further validating our initial conclusions.

The model that we use here is not calibrated to a single tumor or a single virus type. Hence the optimized treatments identified here are not applicable to patients. However, we show that much can be gained by varying OVT scheduling, and it is worthwhile to further explore treatment scheduling in more detail for specific cancers and specific viruses.

## Use of AI tools declaration

The authors declare they have not used Artificial Intelligence (AI) tools in the creation of this article.

## Acknowledgments

We are particularly grateful to discussions with Morgan van Walsum during her NSERC-USRA 2024 scholarship at the University of Alberta. We also thank the Math Bio Journal Club for helpful comments. NM acknowledges the funding support from the University of Alberta. TH is supported through a discovery grant of the Natural Science and Engineering Research Council of Canada (NSERC), RGPIN-2023-04269.

## Appendix

### A.1 Separate compartments for CAR-T cells and general immune cells

Our general immune system consists of both tumour-specific and virus specific immune cells and thus will kill and be stimulated by both cancer cells and the virus. Our suggested extension of (2) is

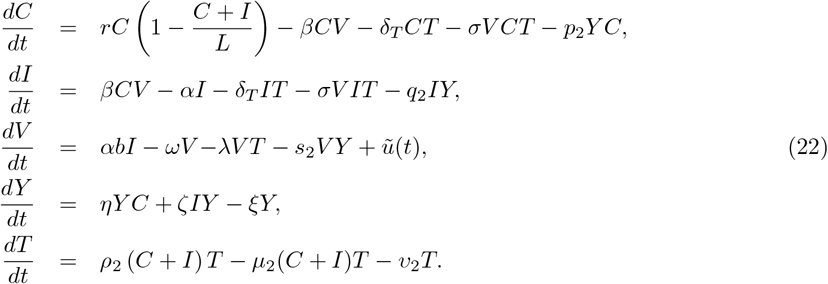

In this model, we consider five cell populations: uninfected cancer cells (*C*), infected cancer cells (*I*), general immune cells (*Y*), CAR-T cells (*T*), and free viruses (*V*). The first four terms in equation one are similar to those in model (20), with the fifth term accounting for the killing of uninfected cancer cells by general immune cells at a rate of *p*_2_. Similarly, in equation two, all terms except the fifth remain the same as in Equation (20), while the last term represents the killing of infected cells by immune cells at a rate of *q*_2_. In equation three, the first two terms describe the effective replication of the virus and its clearance, respectively, while the third term accounts for the elimination of the OV by general immune cells at a rate of *s*_2_. As before *ũ*(*t*) is the dimensional control term and has a similar definition as (16) after non-dimensionalization. Equation four remains identical to the immune cell equation in the base model (3). Finally, in equation five, the first term represents the stimulation and proliferation of CAR-T cells upon interaction with both infected and uninfected cancer cells at a rate of *ρ*_2_. The second term denotes the inactivation of CAR-T cells by both tumor cell types at a rate of *µ*_2_, while the last term represents their natural death at a rate of *υ*_2_.

We see these interactions between the elements in equation (22) more clearly in Figure 10. In this model, the similar parameters as in equation 20 have the same values as in Table 4. The rest of parameters values are given in Table 7. We set the initial conditions to be

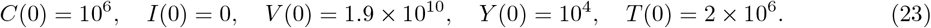

**Table 7:**
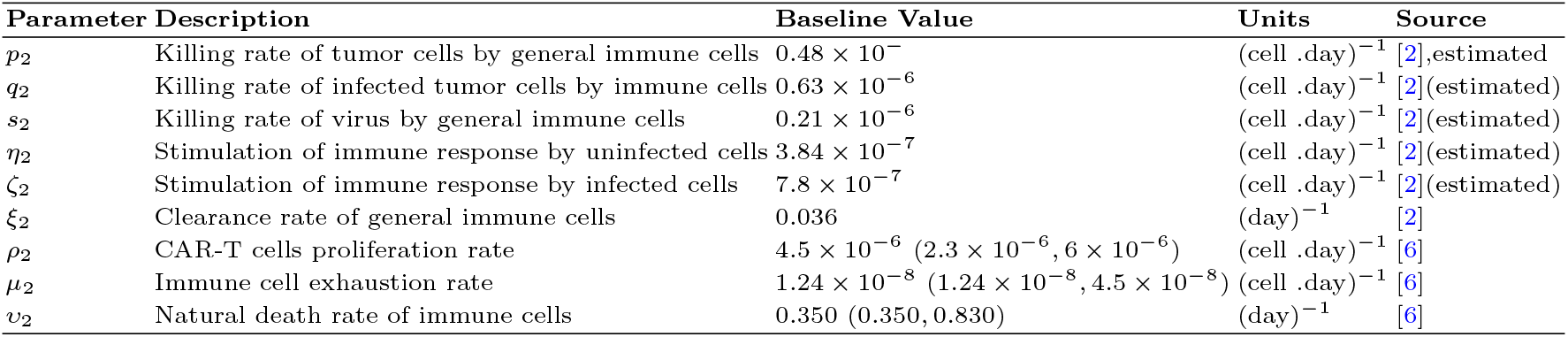
Genreal immune cells and CAR-T cells model parameters, and their baseline values.

**Figure 10:**
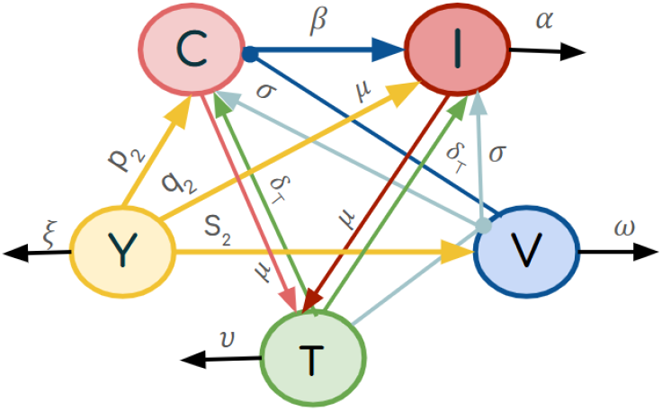
The blue arrows represent the viral infection of cancer cells at a rate *β*. The light cyan arrows indicate the virus-induced cancer cell killing of the CAR-T cells at the rate *σ*, while the orange arrows illustrate their direct killing of the cancer cells at the rate *δ*_*T*_ . The pink and red arrows denote the CAR-T cell death by the cancer cells at the rate *µ*. The interaction of general immune cells with uninfected cancer cell, infected cancer cells, and the virus is denoted in green arrows which the immune cells kill them at the rate *p*_2_, *q*_2_, and *s*_2_ respectively. All black arrows show cell death and virus degradation.

For choosing parameter values, since we are considering the innate immune system, we use the parameters from Al-Tuwairqi et al. [2] in this model as well and for the CAR-T therapy related parameters, we use the values from Barros et al. [6]. We again begin by applying CAR-T cell therapy alone. The simulation results are shown in Figure 11.

**Figure 11:**
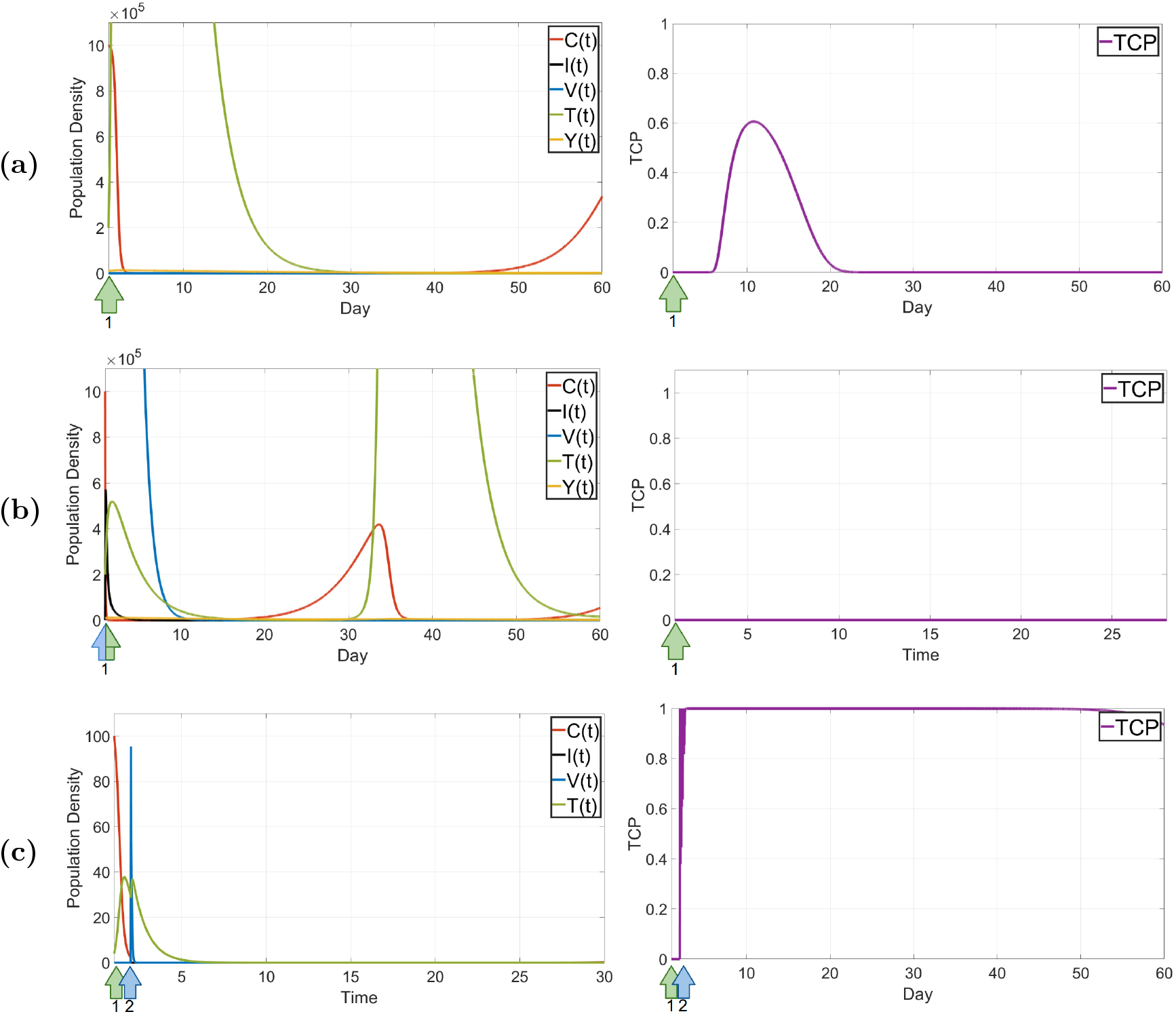
Dynamics of the cancer cells, immune cells, and CAR-T cells. **(a)** No virus and injection of CAR-T cells on day one. **(b)** CAR-T cells and virus both on day one. **(c)** An optimized schedule with virus injection delayed to day two.

### A.2 Model with virus and cancer specific immune cells

Our next step to extend our results is making the immune response more specific. We assume that the T-cells are divided into two groups that primarily target either viral antigens or tumor antigens [37]. Thus, model (2) is extended to:

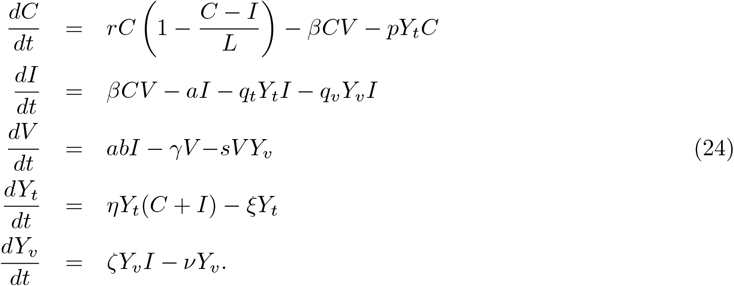

The model now consists of five cell populations: the uninfected tumor cells (*C*), tumor cells infected with OVs (*I*) and tumor and virus specific T cells (*Y*_*t*_) and *Y*_*v*_ respectively, and an OV population (*V*). Same as the base model (2), first term in equation one denotes the logistic growth of the uninfected cell in the absence of the treatment and the second term accounts for the the infection of the cancer cells. The third term in this equation is describing the killing of the cancer cells by the tumor specific immune cells at the rate of *p*. In the second equation the first and the second term are similar the the base model describing the increase in the population of the infected cells and the death of such cells in the rate *a* respectively. The infected cell are marked by the virus so as we see the next two terms we have the death of the infected cells at the rate *q*_*t*_ by the tumor specific immune cells in the third term and also in fourth term their death at the rate *q*_*v*_ by the virus-specific T-cells. In the first term of equation three we have the effective replication of the virus by bursting out of the infected cell at rate *b* and their natural clearance at rate *γ* in the second term. The viruses are killed by the virus-specific immune cells at the rate *s* as described in the last term of equation three. The tumor-specific immune cells are stimulated by both infected and uninfected cancer cells, the first term of equation four denotes this stimulation at the rate *η*, the second term accounts for the clearance of this type of immune cells at the rate *ξ*. In the first term of the last equation we the stimulation of the virus-specific immune cells at the rate *ζ* and the very last term describes their death the rate *ν*. We did not consider any clearance of the immune cells caused by the tumour cells. This was inspired by the model studied in [41]. In [41], K. Storey et al. incorporated both innate and adaptive immune responses in their virotherapy model. They further divided adaptive immune cells into antitumor and antiviral categories.

We see how the cancer cells, virus and the immune cells interact with each other in Figure 12. We simulate the behavior of this model (Figures not shown), and the parameter values are given in Table (8). We assume that at time t = 0 the tumor is at its maximum carrying capacity, with no infected cells. We also assume there is no virus specific immune cell before the injection as they get stimulated once the virus gets administered to the patient. The initial conditions of the system are

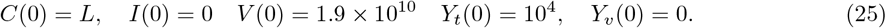

**Table 8:** Immune specific model parameters, and their baseline values.

| Parameter | Description | Baseline Value | Units | Source |
| --- | --- | --- | --- | --- |
| $r$ | Growth rate of uninfected tumor cells | 0.3 | (day)<sup>-1</sup> | [4] |
| $L$ | Tumor carrying capacity | $10^6(10^6 - 10^8)$ | cells | [4] |
| $\beta$ | Infection rate of uninfected tumor cells by free viruses | $1.5 \times 10^{-9}$ | (PFU .day)<sup>-1</sup> | [4] |
| $p$ | Killing rate of tumor cells by tumor-specific immune cells | $0.48 \times 10^{-6}$ | (cell .day)<sup>-1</sup> | [2](estimated) |
| $r$ | Death rate of infected tumor cells | 1 | (day)<sup>-1</sup> | [4] |
| $q_t$ | Killing rate of infected tumor cells by tumor-specific immune cells | $0.63 \times 10^{-6}$ | (cell .day)<sup>-1</sup> | [2](estimated) |
| $q_v$ | Killing rate of infected tumor cells by virus-specific immune cells | $0.63 \times 10^{-6}$ | (cell .day)<sup>-1</sup> | [2](estimated) |
| $b$ | Viruses released by lysed infected tumor cell | 3500 | (PFU)(cell)<sup>-1</sup> | [4] |
| $\gamma$ | Virus-induced immune response rate | 4 | (day)<sup>-1</sup> | [4] |
| $s$ | Killing rate of virus by virus-specific immune cells | $0.21 \times 10^{-6}$ | (cell .day)<sup>-1</sup> | [2](estimated) |
| $\eta$ | Stimulation of immune response by uninfected cells | $3.84 \times 10^{-7}$ | (cell .day)<sup>-1</sup> | [2](estimated) |
| $\zeta$ | Stimulation of immune response by infected cells | $7.8 \times 10^{-7}$ | (cell .day)<sup>-1</sup> | [2](estimated) |
| $\xi$ | clearance rate of tumour-specific immune cells | 0.009 | (day)<sup>-1</sup> | [22] |
| $\nu$ | clearance rate of virus-specific immune cells | 0.1329 | (day)<sup>-1</sup> | [22] |

**Figure 12:**
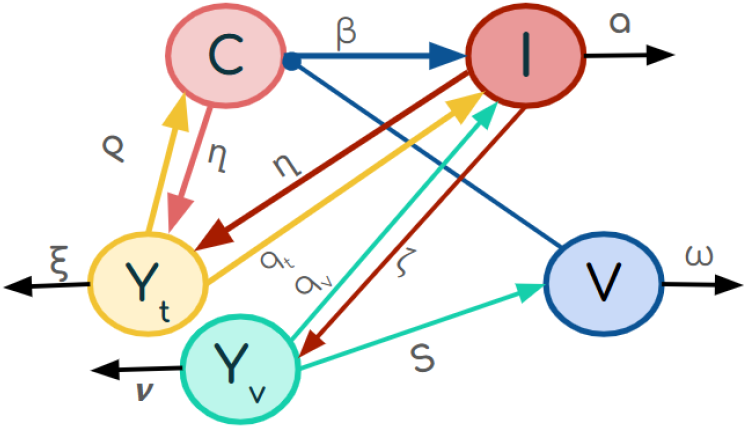
Interactions of the cancer cells, the oncolytic virus and the virus and tumour specific immune cells.

Parameter values were chosen similar to the model (2). We had to stick with the values we had from Al-Twairqi et al. [2] for the immune stimulation related parameter values while for the clearance rate we could use a more specific values taken from Mahasa et al. [22]. The results are summarized in Table 6.

